## Supplementary materials for "Statistical integration of multi-omics and drug screening data from cell lines"

**1** Dept. Data science & Biostatistics, UMC Utrecht, Utrecht, Netherlands
**2** Department of Neurology, Hannover Medical School, Hannover, Germany
**3** Roche Institute for Translational Bioengineering, Basel, Switzerland
**4** Max Planck Institute of Molecular Cell Biology and Genetics, Dresden, Germany
**5** Department of Neurology, Ludwig-Maximilians-Universität, Munich, Germany
**6** German Center for Neurodegenerative Diseases, Munich, Germany
**7** Dept. of Mathematics, Radboud University, Nijmegen, Netherlands

†These authors contributed equally to this work.
‡These authors also contributed equally to this work.
*

### 1 Introduction

This document is a supplement to the main article “Statistical integration of multi-omics data identifies potential therapeutic targets for MSA”. It is structured into two parts.

The first part, Section 2, contains the mathematical details of POPLS-DA and corresponding proofs. Also, the maximum likelihood estimation algorithm is presented. We discuss a strategy to select the number of joint components and the number of relevant genes. We additionally discuss how to interpret the POPLS-DA parameters.

The second part contains additional tables and figures for the multi-omics and bioinformatics analyses. We show scree plots for selecting the number of joint components and number of relevant genes. We perform a permutation test to assess whether the top 200 genes can significantly distinguish the two experimental groups. A table of these relevant genes is presented, as well as the GO enrichment clusters for those genes that were targeted by a drug.

### 2 Methods

For a joint analysis of the transcriptomics and proteomics datasets, we propose probabilistic orthogonal partial least squares discriminant analysis (POPLS-DA).

#### 2.1 Mathematical model of POPLS-DA

As in the main article, $x_t$ and $x_p$ are random vectors of size $d$ representing the transcriptomics and proteomics data. Further, $y_t$ and $y_p$ are univariate random variables representing dummy variables for the experimental conditions (one for cells overexpressing $\alpha$Syn, and zero for the controls). In the POPLS-DA model, the omics data and the experimental conditions are linked via latent variables $u_t$ and $u_p$ of size $r$, where $r$ is much smaller than the data dimensions. Specific components $v_t$ and $v_p$ are added to the model to capture variation of the omics data that doesn’t play a role in discriminating the two experimental groups. The numbers of specific components are $r_t$ and $r_p$, respectively. Residual variation is modeled by including noise terms $e_t$ and $e_p$ for the transcriptomics and proteomics datasets, and $\epsilon_t$ and $\epsilon_p$ for the random variable representing the two conditions. The joint and specific latent variables $u_t$, $u_p$, $v_t$ and $v_p$ are assumed to be standard normally distributed. The residual terms $e_t$, $e_p$, $\epsilon_t$ and $\epsilon_p$ are normally distributed with zero mean and (co)variance (matrix) $\sigma^2_{e_t} I_d$, $\sigma^2_{e_p} I_d$, $\sigma^2_{\epsilon_t}$, and $\sigma^2_{\epsilon_p}$.

The mathematical model can be written as

$$\begin{aligned} x_t &= u_t W^{\mathrm{T}} + v_t P_t^{\mathrm{T}} + e_t \qquad & x_p &= u_p W^{\mathrm{T}} + v_p P_p^{\mathrm{T}} + e_p, \\ y_t &= u_t \beta + \epsilon_t & y_p &= u_p \beta + \epsilon_p. \end{aligned} \tag{1}$$

The matrices $W$ of size $d \times r$, $P_t$ of size $d \times r_t$ and $P_p$ of size $d \times r_p$ contain the joint and specific loadings for each gene. The vector $\beta$ contains the regression coefficients of $y_t$ and $y_p$ on the joint components $u_t$ and $u_p$.

In the POPLS-DA model, $(x_t, y_t)$ is normally distributed with zero mean and covariance matrix

$$\Sigma_t = \begin{bmatrix} WW^{\mathrm{T}} + P_t P_t^{\mathrm{T}} + \sigma^2_{e_t} I_d & W\beta \\ \beta W^{\mathrm{T}} & \beta^{\mathrm{T}}\beta + \sigma^2_{\epsilon_t} \end{bmatrix}. \tag{2}$$

Analogously, $(x_p, y_p)$ is normally distributed with zero mean and covariance matrix denoted by $\Sigma_p$. The set of all parameters is denoted by $\theta = \{W, \beta, P_t, P_p, \sigma_{e_t}, \sigma_{e_p}, \sigma_{\epsilon_t}, \sigma_{\epsilon_p}\}$.

#### 2.2 Estimation with maximum likelihood

The data from all samples are collected in $X_t$, $X_p$ $Y_t$, and $Y_p$. Let $X = [X_t^{\mathrm{T}}, X_p^{\mathrm{T}}]^{\mathrm{T}}$ and $Y = [Y_t^{\mathrm{T}}, Y_p^{\mathrm{T}}]^{\mathrm{T}}$ be the datasets stacked across the rows. We propose to optimize the POPLS-DA likelihood by implementing an expectation-maximization (EM) algorithm. The log likelihood is given by

$$l(\theta|X, Y) = f((X_t, Y_t), \mu = 0, \Sigma = \Sigma_t) \cdot f((X_p, y_p), \mu = 0, \Sigma = \Sigma_p) \tag{3}$$

where $f$ is a multivariate normal density function with the given mean and covariance matrix.

We propose to optimize the likelihood using EM, where the complete data likelihood can be written (with abuse of notation) as

$$l_c(\theta) = f(X_t|U_t, V_t) f(Y_t|U_t) f(U_t) f(V_t) \cdot f(X_p|U_p, V_p) f(Y_p|U_p) f(U_p) f(V_p). \tag{4}$$

In an EM algorithm, the conditional expectation of $l_c(\theta)$ given $X$ and $Y$ is optimized. Note that the optimizations can be decoupled and performed per term. For example, the first term yields the optimization problem,

$$\max_\theta \mathbb{E}[\log f(X_t|U_t, V_t)|X_t, Y_t] + \mathbb{E}[\log f(X_p|U_p, V_p)|X_p, Y_p]. \tag{5}$$

With respect to $W$, this becomes

$$\min_W \mathbb{E}[||X_t - U_t W^{\mathrm{T}} - V_t P_t^{\mathrm{T}}||_F^2 | X_t, Y_t] + \mathbb{E}[||X_p - U_p W^{\mathrm{T}} - V_p P_p^{\mathrm{T}}||_F^2 | X_p, Y_p]. \tag{6}$$

In the expectation step, we calculate the conditional expectations of the latent variables. We focus on the $x_t$ terms. First, note that we can rewrite the POPLS-DA model as

$$(x_t, y_t) = (u_t, v_t)\Gamma^{\mathrm{T}} + (e_t, \epsilon_t), \tag{7}$$

with $\Gamma_t = \begin{pmatrix} W & P \\ \beta^{\mathrm{T}} & 0 \end{pmatrix}$. Using the rules for conditional expectations of normal distributions (e.g. see [1]) , we get

$$\begin{aligned} \mathbb{E}[(u_t, v_t)|x_t, y_t] &= (x_t, y_t)\Sigma^{-1}_{(e_s,\epsilon_s)}\Gamma\tilde{\Sigma}_{u_t,v_t}, \\ \mathrm{Var}[(u_t, v_t)|x_t, y_t] &= \tilde{\Sigma}_{u_t,v_t} \end{aligned} \tag{8}$$

where $\tilde{\Sigma}_{u_t,v_t} = (I - \Gamma^{\mathrm{T}}\Sigma^{-1}_{(e_s,\epsilon_s)}\Gamma)$. By replacing $t$ with $p$, the conditional expectations given $(x_p, y_p)$ are obtained. With these expressions, the objective function (6) can be optimized.

In the maximization step for $W$, we solve the optimization problem (6). Note that both terms depend on $W$. Taking the derivative with respect to $W$ and equating to zero yields

$$\begin{aligned} X_t^{\mathrm{T}}\mathbb{E}[U_t|X_t, Y_t] - P_t\mathbb{E}[U_t^{\mathrm{T}}V_t|X_t, Y_t] + X_p^{\mathrm{T}}\mathbb{E}[U_p|X_p, Y_p] - P_p\mathbb{E}[U_p^{\mathrm{T}}V_p|X_p, Y_p] = \\ W\{\mathbb{E}[U_t^{\mathrm{T}}U_t|X_t, Y_t] + \mathbb{E}[U_p^{\mathrm{T}}U_p|X_p, Y_p]\} \end{aligned} \tag{9}$$

The expected values are taken from the expectation step. This yields the maimization step for $W$, which is given below. Optimizers for the other terms of the complete likelihood are obtained similarly. The complete EM algorithm for POPLS-DA is given by the following iterative scheme in $k$, starting with initial values for $k = 0$ and $s = t, p$,

$$\begin{aligned} W^{k+1} &= (\sum_{s=t,p} X_s^{\mathrm{T}}\mathbb{E}_k[U_s] - P_s^k\mathbb{E}_k[V_s]^{\mathrm{T}}\mathbb{E}_k[U_s])(\sum_{s=t,p}\mathbb{E}_k[U_s^{\mathrm{T}}U_s])^{-1} \\ \beta^{k+1} &= (\sum_{s=t,p} Y_s^{\mathrm{T}}\mathbb{E}_k[U_s])(\sum_{s=t,p}\mathbb{E}_k[U_s^{\mathrm{T}}U_s])^{-1} \\ P_s^{k+1} &= (X_s^{\mathrm{T}}\mathbb{E}_k[V_s] - W^{k+1}\mathbb{E}_k[U_s]^{\mathrm{T}}\mathbb{E}_k[V_s])(\mathbb{E}_k[V_s^{\mathrm{T}}V_s])^{-1} \\ (\sigma^2_{e_s})^{k+1} &= (N_s d)^{-1}\mathbb{E}_k[E_s^{\mathrm{T}}E_s] \\ (\sigma^2_{\epsilon_s})^{k+1} &= (N_s)^{-1}\mathbb{E}_k[\mathcal{E}_s^{\mathrm{T}}\mathcal{E}_s] \end{aligned} \tag{10}$$

In this scheme, $\mathbb{E}_k[\cdot] = \mathbb{E}[\cdot|X, Y, \theta^k]$.

#### 2.3 Interpretation of POPLS-DA.

For sake of brevity, we will drop the subscript $t$ and $p$ in this paragraph. In the article, $x$ in Equation (1) represents the transcriptomics and proteomics data. The experimental conditions for each omics dataset are indicated by $y$. POPLS-DA models the relationship between all omics data and conditions simultaneously in terms of the joint latent variables $u$. The loadings $W$ represent the projection of the omics data onto the $u$. In our model, the loading weight for gene $j$ in joint principal component $k$ is given by $w_{j,k}$. A larger weight indicates a larger relative contribution to that joint component. Furthermore, if $w_{j,k}$ and $w_{j',k}$ have the same sign, the corresponding genes $j$ and $j'$ are positively correlated within component $k$. The number $(W\beta)_j = \sum_k w_{j,k}\beta_k$ is the coefficient indicating the relationship between gene $j$ with the experimental groups. The vector $W\beta$ can be interpreted as the covariance of $y$ with all omics data $x$. The latent variable $u_{i,k}$ indicates the position of data point $x_i$ in the joint component $k$. Two subjects $i$ and $i'$ are similarly positioned if $u_{i,k}$ and $u_{i',k}$ are similar numbers. The interpretation of $v$ and $P$ goes analogously.

#### 2.4 Dimension selection.

The POPLS-DA model is formulated conditional on the number of joint and specific components, $r$ and $r_x$, respectively. To estimate the number of components and the number of variables to investigate, we consider scree plots [2]. For the number of components, we calculate the eigenvalues of the stacked dataset $X = [X_1^{\mathrm{T}}, \ldots, X_s^{\mathrm{T}}]^{\mathrm{T}}$ and inspect the corresponding scree plot. To determine the number of variables per component to investigate, we calculate the estimated effect size $W\beta$ of each variable and use the "elbow" criterion on the sorted effect sizes (as in a scree plot [2]).

The proportions of variance explained by the joint and specific parts are calculated by dividing the trace of the corresponding covariance matrices by the trace of the modeled covariance matrix. In particular, the proportion of the variance of $x_t$ explained by the joint part can be calculated by $\mathrm{tr}W^{\mathrm{T}}W$ divided by $\mathrm{tr}W^{\mathrm{T}}W + \mathrm{tr}P_t^{\mathrm{T}}P_t + d\sigma_{e_t}^2$.

### 3 Results

In this section, we present the supplementary figures and tables corresponding to the data analysis in the main article.

Scree plots of the eigenvalues of transcriptomics and proteomics datasets are shown in Figure 2, together with a scree plot of the effect sizes $W\beta$. Based on visually assessing where a plateau occurs, two joint and two specific components were selected. Furthermore, based on the visual assessment of the occurrence of an elbow, 200 genes were retained for interactome and functional analyses. These genes are given in Table 1.

We performed a permutation test where we randomly shuffled the case-control label for each sample. We then apply POPLS-DA again and calculate the overall accuracy (number of samples correctly classified based on the top 200 genes). The number of permutations was 400. In one permutation round, POPLS-DA achieved a perfect classification, corresponding with a p-value of 0.0025 (standard error: 0.0025). The median overall accuracy was 41.7% (equivalent to 10 out of 24 samples correct classifications).

The GO enrichment clusters for the subset of genes targeted by a drug are shown in Table **??**. The list of drugs with their direct neighbors are shown in Table **??**.

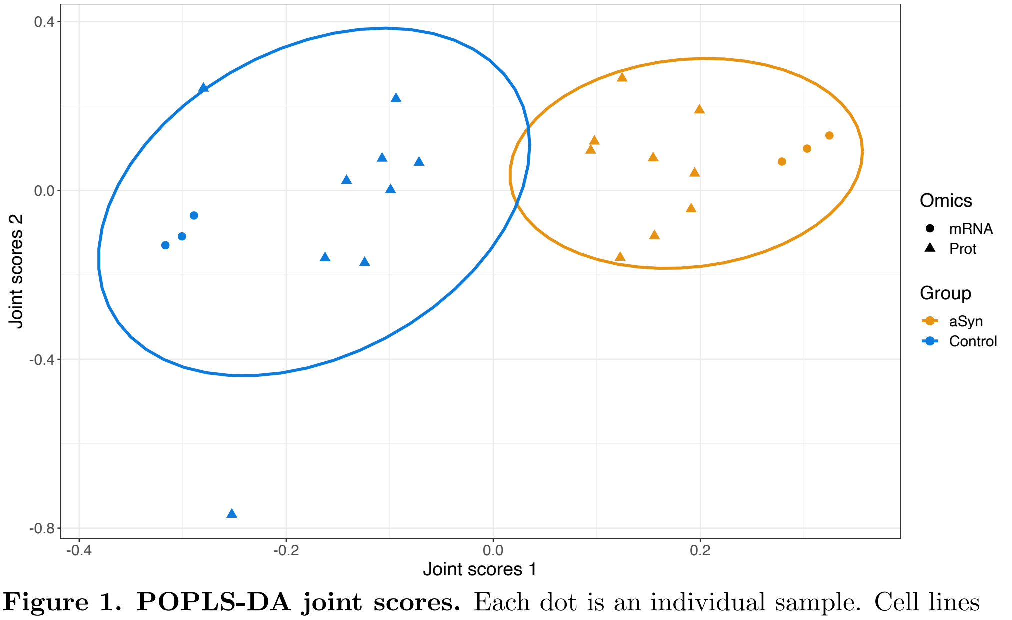


**Figure 1. POPLS-DA joint scores.** Each dot is an individual sample. Cell lines overexpressing $\alpha$-synuclein are indicated by an yellow color, cell lines overexpressing the control protein are colored blue. A 90% confidence ellipse is added based on a $t$ distribution.

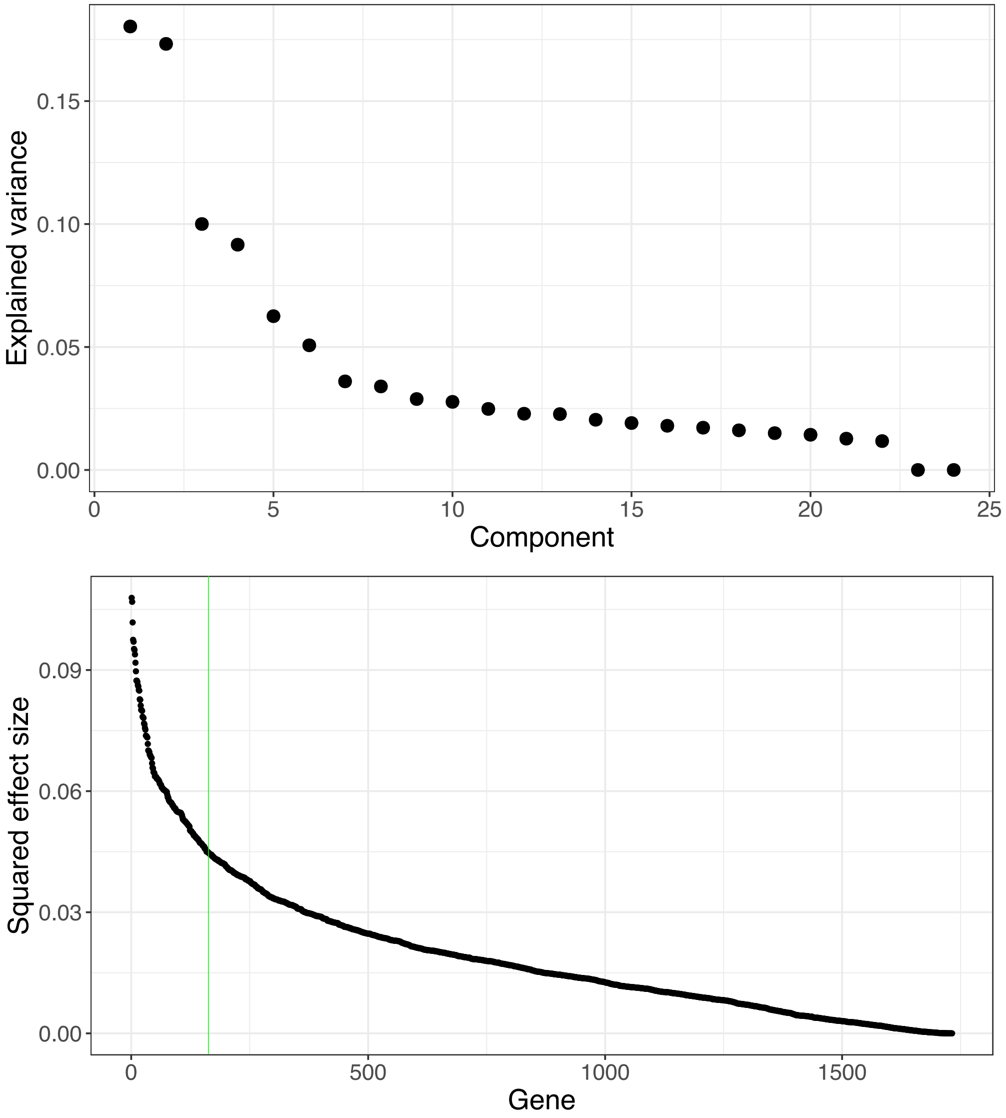


**Figure 2. Scree plots for POPLS-DA**. The number of components or genes is selected by visually assessing where a plateau or 'elbow' curve occurs. The *upper panel* shows the squared singular values of the stacked transcriptomics and proteomics data. The numbers are rescaled to sum up to one. The *lower panel* shows the sorted squared effect size per gene given by the squared elements of $W\beta$.

| weight | SYMBOL | weight | SYMBOL | weight | SYMBOL | weight | SYMBOL |
|---|---|---|---|---|---|---|---|
| 0.338 | MTHFD2 | 0.264 | ESYT1 | 0.242 | DDX3X | 0.227 | AATF |
| 0.33 | CBS | 0.264 | SLC4A7 | 0.242 | MXRA7 | 0.227 | ARSA |
| 0.328 | SCG2 | 0.264 | PLD3 | 0.241 | RPL11 | 0.227 | UFD1L |
| 0.326 | TFRC | 0.262 | GPC6 | 0.24 | PHGDH | 0.227 | MYO5A |
| 0.322 | NES | 0.262 | ALDH6A1 | 0.24 | P2RX3 | 0.227 | RP2 |
| 0.322 | SLC3A2 | 0.262 | MYCBP2 | 0.24 | ARHGAP17 | 0.226 | RPL6 |
| 0.322 | SCARB2 | 0.262 | RRM1 | 0.24 | EEF2 | 0.226 | WARS |
| 0.322 | FAT1 | 0.261 | MAPK8IP3 | 0.239 | SPTLC2 | 0.226 | REM2 |
| 0.318 | SLC7A5 | 0.261 | SARM1 | 0.239 | NEDD4L | 0.226 | ACBD3 |
| 0.318 | POLDIP3 | 0.261 | RCN2 | 0.239 | TJP1 | 0.226 | PDIA6 |
| 0.313 | NRP2 | 0.261 | PEG10 | 0.239 | PON2 | 0.225 | ENDOD1 |
| 0.312 | ASNS | 0.261 | P4HB | 0.238 | HSPA5 | 0.225 | SND1 |
| 0.312 | AIMP2 | 0.261 | SEC61A1 | 0.238 | RPL38 | 0.225 | NAV1 |
| 0.306 | TNC | 0.26 | BCAT1 | 0.238 | RPSA | 0.225 | PYCR1 |
| 0.306 | CNTN2 | 0.259 | ZYX | 0.238 | HYOU1 | 0.224 | LAMP1 |
| 0.302 | FABP3 | 0.258 | DDX17 | 0.238 | PSME2 | 0.224 | TOR1B |
| 0.301 | RPS4X | 0.258 | GNS | 0.237 | MARS | 0.224 | SMPD1 |
| 0.301 | SYT1 | 0.256 | STX12 | 0.236 | RRAS2 | 0.224 | YBX1 |
| 0.298 | STXBP1 | 0.256 | ERGIC1 | 0.236 | HSPB1 | 0.223 | RAD21 |
| 0.297 | SRL | 0.256 | ABCB6 | 0.235 | ALDH9A1 | 0.223 | CARS |
| 0.297 | PAFAH1B1 | 0.256 | HPDL | 0.235 | VAV2 | 0.223 | SEMA6D |
| 0.296 | PSAT1 | 0.255 | CKMT1B | 0.235 | GAA | 0.223 | ADAR |
| 0.296 | KIF5C | 0.254 | NEFM | 0.234 | NUDT21 | 0.223 | EEF1B2 |
| 0.294 | BRD3 | 0.253 | RPAP3 | 0.234 | ARID3B | 0.223 | CSTF3 |
| 0.293 | CNTNAP1 | 0.252 | PRKAR2A | 0.234 | LUC7L2 | 0.223 | SLC2A1 |
| 0.287 | EEF1A2 | 0.251 | ESD | 0.234 | KATNB1 | 0.223 | POLR2A |
| 0.285 | OSBP | 0.251 | ENO3 | 0.234 | ADD1 | 0.222 | C11orf54 |
| 0.284 | VIM | 0.251 | IQSEC1 | 0.234 | CALU | 0.222 | RPS7 |
| 0.283 | ACAT1 | 0.25 | ATP6V1C1 | 0.234 | VCL | 0.222 | HMGCR |
| 0.283 | IGF2R | 0.25 | GATAD2A | 0.234 | AP2A2 | 0.222 | RPS3 |
| 0.282 | SHMT2 | 0.249 | VAT1 | 0.233 | GLUD1 | 0.222 | CLCN6 |
| 0.282 | DNAJB11 | 0.249 | USP9X | 0.233 | HDGFRP3 | 0.222 | GLDC |
| 0.282 | NQO1 | 0.248 | RPS24 | 0.233 | YTHDC1 | 0.222 | RPL4 |
| 0.281 | RPL7A | 0.248 | SDF4 | 0.233 | DICER1 | 0.221 | EIF4B |
| 0.28 | CCDC50 | 0.248 | YARS | 0.232 | RPS21 | 0.221 | COL6A1 |
| 0.277 | ECEL1 | 0.248 | TAF15 | 0.232 | CYB5A | 0.221 | RCC1 |
| 0.277 | ALDH7A1 | 0.246 | PDLIM3 | 0.232 | GBA2 | 0.221 | CADM4 |
| 0.277 | GARS | 0.246 | ACTC1 | 0.231 | NDUFA11 | 0.221 | GOLM1 |
| 0.275 | SPTBN1 | 0.246 | FAR1 | 0.231 | SMARCA1 | 0.221 | AARS |
| 0.275 | MAP2K6 | 0.246 | PDCD4 | 0.231 | XPO5 | 0.22 | THNSL1 |
| 0.27 | MDGA1 | 0.245 | HAX1 | 0.231 | NCBP1 | 0.22 | FAM171A1 |
| 0.269 | RAB10 | 0.245 | RAPGEF6 | 0.23 | TARS | 0.22 | SLC7A6 |
| 0.267 | RABGAP1 | 0.245 | ANK2 | 0.23 | MAP1A | 0.22 | CTSC |
| 0.267 | FYN | 0.244 | DSG2 | 0.23 | MUT | 0.22 | SLC9A3R1 |
| 0.267 | RCN1 | 0.244 | RPL8 | 0.229 | SYT11 | 0.22 | SNRNP200 |
| 0.266 | EPB41L5 | 0.244 | SMC3 | 0.229 | GRIPAP1 | 0.219 | CLIC1 |
| 0.265 | EIF5 | 0.243 | OXCT1 | 0.229 | APLP2 | 0.219 | DPM1 |
| 0.265 | LGALS3BP | 0.242 | FAHD2A | 0.229 | NAPB | 0.219 | HDGF |
| 0.265 | SLC1A4 | 0.242 | FKBP2 | 0.229 | NDUFA13 | 0.219 | AKAP12 |
| 0.265 | RFX3 | 0.242 | ATP2C1 | 0.228 | SPIN1 | 0.219 | PSIP1 |

**Table 1. Top 200 genes in the joint transcriptomics-proteomics part.** The weights are estimated with POPLS-DA, and indicate the contribution of each gene towards the classification.

| Rank | Term | Ontology | Genes in set |
|---|---|---|---|
| 1 | extracellular exosome | CC | 7.54e-22 |
| 2 | extracellular vesicle | CC | 7.78e-22 |
| 3 | extracellular membrane-bounded organelle | CC | 7.94e-22 |
| 4 | extracellular organelle | CC | 7.94e-22 |
| 5 | cytoplasm | CC | 2.39e-17 |
| 6 | vesicle | CC | 2.69e-16 |
| 7 | extracellular space | CC | 1.49e-15 |
| 8 | cell junction | CC | 1.05e-14 |
| 9 | extracellular region | CC | 3.19e-13 |
| 10 | translation | BP | 5.82e-12 |
| 11 | amide biosynthetic process | BP | 7.94e-12 |
| 12 | peptide biosynthetic process | BP | 1.33e-11 |
| 13 | melanosome | CC | 2.79e-11 |
| 14 | pigment granule | CC | 2.79e-11 |
| 15 | RNA binding | MF | 7.63e-11 |
| 16 | cytoplasmic translation | BP | 1.09e-10 |
| 17 | developmental process | BP | 1.33e-10 |
| 18 | cellular amide metabolic process | BP | 1.59e-10 |
| 19 | focal adhesion | CC | 2.08e-10 |
| 20 | cell-substrate junction | CC | 2.98e-10 |

**Table 2. Gene ontology enrichment analysis of drug targeted genes.** A subset of the top 200 genes, consisting of 116 genes that are (indirectly) targeted by an FDA approved drug compound, were used to perform GO enrichment analysis. The twenty most significant terms are shown.

| Drug | Genes |
|---|---|
| Ajmaline | SPTBN1, ACTC1, ANK2, DSG2, SPTLC2, NEDD4L, NAV1 |
| Amiodarone | CBS, SCG2, TFRC, NES, SCARB2, FAT1, TNC, FABP3, SYT1, EEF1A2, VIM, IGF2R, NQO1, FYN, RCN1, LGALS3BP, P4HB, GATAD2A, ACTC1, PDCD4, HAX1, EEF2, TJP1, PON2, HSPA5, RPSA, HSPB1, ALDH9A1, ADD1, CALU, DICER1, GBA2, NCBP1, APLP2, WARS, NAV1, PYCR1, LAMP1, YBX1, SLC2A1, HMGCR, CTSC, SLC9A3R1, CLIC1 |
| Amlodipine | SCARB2, SYT1, STXBP1, NQO1, ESD, PDLIM3, ACTC1, HAX1, ANK2, SPTLC2, HSPA5, GAA, ADD1, GBA2, ARSA, REM2, NAV1, LAMP1, YBX1, SLC2A1, HMGCR |
| Astemizole | CBS, NES, CNTN2, STXBP1, KIF5C, NQO1, MDGA1, FYN, P4HB, NEFM, HAX1, ANK2, DSG2, EEF2, NEDD4L, HSPA5, ADD1, GBA2, NDUFA11, SYT11, APLP2, AATF, NAV1, YBX1, SLC2A1, HMGCR |
| Benazepril | MTHFD2, CBS, SLC3A2, SLC7A5, SHMT2, NQO1, SLC1A4, ABCB6, DSG2, MARS, GAA, ADD1, MUT, SLC2A1, HMGCR, CLCN6, GLDC, SLC7A6 |
| Bepridil | POLDIP3, SYT1, RPS4X, STXBP1, VIM, IGF2R, NQO1, RCN2, PEG10, SEC61A1, ABCB6, ACTC1, HAX1, ANK2, DSG2, RPL8, DDX3X, RPL11, EEF2, NEDD4L, HSPA5, RPL38, ADD1, AP2A2, RPS21, GBA2, SYT11, MYO5A, REM2, NAV1, LAMP1, YBX1, SLC2A1, RPS7, HMGCR, RPS3 |
| Pentoxyverine | ACAT1, ANK2, DSG2, P2RX3, NEDD4L, HSPA5, NAV1, YBX1, HMGCR, SLC9A3R1 |
| Clemastine | NQO1 |
| Dexibuprofen | CBS, SCG2, TFRC, NES, SLC3A2, SCARB2, FAT1, SLC7A5, TNC, FABP3, SYT1, EEF1A2, VIM, IGF2R, NQO1, FYN, RCN1, LGALS3BP, SLC1A4, SLC4A7, P4HB, SEC61A1, GNS, ABCB6, PRKAR2A, GATAD2A, VAT1, SDF4, ACTC1, PDCD4, HAX1, DSG2, RPL8, SMC3, EEF2, NEDD4L, TJP1, PON2, HSPA5, RPSA, PSME2, HSPB1, ADD1, CALU, AP2A2, DICER1, GBA2, SYT11, APLP2, WARS, PYCR1, LAMP1, YBX1, SLC2A1, HMGCR, RPS3, SLC7A6, CTSC, SLC9A3R1, CLIC1 |
| Dicyclomine | TFRC, SLC3A2, SCARB2, SYT1, IGF2R, FKBP2, AP2A2, SYT11 |
| Dipyridamole | DDX17, PRKAR2A, ESD, PDLIM3, HAX1, RPL11, SPTLC2, HSPA5, ALDH9A1, DICER1, GBA2, YBX1, SLC2A1, HMGCR, AARS, SLC9A3R1 |
| Doxazosin | NQO1, HAX1, HSPA5, CALU, GBA2, YBX1, SLC2A1, HMGCR |
| Dyclonine | SPTBN1, ANK2, P2RX3, NAV1 |
| Flunarizine | POLDIP3, SYT1, NQO1, RCN2, SEC61A1, ACTC1, ANK2, DDX3X, EEF2, ADD1, CALU, MYO5A, NAV1, HMGCR |
| Guanfacine | SLC7A5, IGF2R, NQO1, ALDH9A1, HMGCR |
| Ifenprodil | CNTN2, SYT1, STXBP1, SPTBN1, FYN, ANK2, APLP2 |
| Imatinib | CBS, SCG2, TFRC, NES, NRP2, TNC, FABP3, SYT1, EEF1A2, VIM, IGF2R, NQO1, MAP2K6, FYN, RCN1, LGALS3BP, P4HB, ACTC1, HAX1, P2RX3, EEF2, SPTLC2, NEDD4L, TJP1, PON2, HSPA5, RPSA, HSPB1, ALDH9A1, VAV2, ADD1, CALU, AP2A2, DICER1, GBA2, APLP2, PYCR1, LAMP1, YBX1, SLC2A1, POLR2A, HMGCR, RPS3, CADM4, CTSC, SLC9A3R1, CLIC1 |
| Lomerizine | NQO1, HAX1, HSPA5, GBA2, YBX1, SLC2A1, HMGCR |
| Nefazodone | SCG2, NQO1, NEFM, HAX1, ANK2, DSG2, NEDD4L, HSPA5, GBA2, MAP1A, NAV1, YBX1, CARS, SLC2A1, HMGCR |
| Pentamidine | SLC3A2, NQO1, YARS, ALDH9A1, ADD1, DICER1 |
| Quinacrine | PAFAH1B1, VIM, IGF2R, NQO1, PLD3, HAX1, HSPA5, ALDH9A1, ADD1, GBA2, YBX1, SLC2A1, HMGCR |
| Reserpine | SCG2, SYT1, PAFAH1B1, IGF2R, EIF5, RRM1, USP9X, HAX1, SMC3, HSPA5, ALDH9A1, ADD1, GBA2, YBX1, RAD21, SLC2A1, HMGCR, SLC9A3R1 |

*Continues on next page*

| Drug | Genes |
|---|---|
| Risperidone | CBS, SCG2, TFRC, NES, FAT1, TNC, FABP3, SYT1, EEF1A2, VIM, IGF2R, NQO1, FYN, RCN1, LGALS3BP, P4HB, NEFM, ACTC1, HAX1, EEF2, SPTLC2, TJP1, PON2, HSPA5, RPSA, HSPB1, ADD1, CALU, DICER1, GBA2, MAP1A, APLP2, PYCR1, LAMP1, YBX1, CARS, SLC2A1, HMGCR, CTSC, SLC9A3R1, CLIC1 |
| Telmisartan | TFRC, NES, SCARB2, FABP3, SYT1, IGF2R, NQO1, GATAD2A, PDCD4, HAX1, TJP1, HSPA5, RPL38, ADD1, AP2A2, DICER1, GBA2, SYT11, WARS, SND1, YBX1, SLC2A1, HMGCR, SLC9A3R1 |
| Triflupromazine | TFRC, SLC3A2, SCARB2, SYT1, IGF2R, HAX1, FKBP2, PON2, HSPA5, RRAS2, AP2A2, GBA2, SYT11, YBX1, SLC2A1, HMGCR |
| Trimipramine | SCG2, FAT1, NQO1, GARS, NEFM, HAX1, P2RX3, HSPA5, CALU, GBA2, MAP1A, YBX1, CARS, SLC2A1, HMGCR |
| Tropisetron | FAT1, GARS, P2RX3 |

**Table 3. List of the drugs and their direct neighbors.** For each validated drug, the genes in the top 200 that interact with any drug target are shown .
